# Generation of uniform-sized spheroids using PDMS-based microwell devices as 3D cancer models for pharmacological profiling of drugs

**DOI:** 10.1101/2025.09.01.673452

**Authors:** Sowjanya Goli, Aditya Teja Guduru, Suman Karadagatla, Errappagari Sreehari, Ira Bhatnagar, Manoj P. Dandekar, Anamika Sharma, Dhiraj Bhatia, Amit Asthana

## Abstract

Cancer researchers now consider spheroids a valuable in vitro model for cancer research and personalized medicine. They are used to studying cancer development, testing drug effectiveness, and potentially guiding treatment decisions for individual patients. Spheroids represent the simplest form of three-dimensional (3D) cellular arrangement and encapsulate the essential tumor microenvironment. These characteristics are crucial for studying processes such as tumor invasion, metastasis, angiogenesis, and cell cycle kinetics. In particular, spheroids excel in chemo-response assays where traditional monolayer cell cultures often fall short. A PDMS microwell device was developed to generate uniform-sized cancer spheroids. This device is user-friendly and capable of producing a large number of spheroids. The device measures 13 mm in diameter (1200 microwells per well if the device has microwells 400 μm in size, and 300 microwells per well if the device has microwells 800 μm in size). It advances 3D cultures by requiring only a small volume of cell culture supplements and is easy to manage. The hydrophobic nature of the PDMS device prevents cells from adhering to it, thereby promoting spheroid formation. Spheroids can be created on microwell devices, and subsequent experiments may either be conducted on the device or transferred to cell culture dishes for additional 3D biological assays. Seeding cells is notably easier compared to other 3D cell culture techniques, and the number of cells in each spheroid can be adjusted according to specific requirements. Overall, the PDMS-based microwell device offers a simple and efficient means to produce large quantities of uniform-sized spheroids for 3D cell culture studies, showcasing high throughput, short generation times, long-term effectiveness, and ease of handling.

## Introduction

Pharmacological profiling employs both in vitro and in-vivo assays to evaluate the potential and safety risks of a lead molecule during the initial stages of drug discovery [1]. Tumor heterogeneity complicates cancer treatment [2], 2D cell cultures frequently do not replicate *in-vivo* responses. [3], leading to incorrect results. Personalized medicine tailors treatments to individual genetic and environmental factors, thereby enhancing effectiveness while minimizing side effects [4]. Personalized approaches also use patient-derived cells for drug screening, offering more relevant preclinical models. In contrast to 2D cultures, 3D models such as spheroids more accurately mimic *in-vivo* environments [5], improving drug penetration and efficacy prediction, making them more reliable for in vitro drug testing [6].

Multicellular spheroids establish gradients reflecting proliferation rates, identifiable as three distinct zones. The outer proliferating zone consists of 4-5 cell layers that imitate regions adjacent to angiogenesis-induced capillaries. The middle quiescent zone houses living cells that cannot proliferate due to restricted oxygen and nutrients. The necrotic core, characterized by hypoxic conditions, resembles tumors lacking a blood supply, typically observed in spheroids larger than 500 μm in size. Investigating cells within spheroids is recognized as 3D cell modeling [7, 8].

There are several methods for performing 3D cell cultures, such as the hanging drop method [9], AggreWell™ plates [10], low retention U bottom plates [11], liquid overlay techniques [12], etc. All these techniques are non-microfluidic [13]. Several recent studies have included microfluidic techniques that help generate multicellular spheroids [14].

The hanging-drop method promotes cell aggregation through gravity but causes oxygen and nutrient scarcity, affecting spheroid physiology and drug testing [15, 16]. Spinner flasks and stirred tanks provide straightforward, economical solutions for producing 3D cell cultures; however, they necessitate substantial media volumes and space, and the stirring speed may cause damage to spheroids [17] or even prevent their formation [18, 19].

The liquid overlay technique, using agarose gel to inhibit cell adhesion [20], encourages spheroid formation[21] but can also induce resistance to therapeutics due to its interaction with the physiology of cells [22]. Non-microfluidic methods like U-bottom[23] and AggreWell™ plates[24] generate spheroids but suffer from issues like non-uniform sizes, labor-intensive handling, and limited throughput, making them less effective for pharmacological profiling [25] compared to microfluidic methods. Despite these limitations, non-microfluidic techniques have been essential in the establishment of this field [26].

Microfluidic techniques have significantly contributed to the advancement of 3D spheroid research by resolving the constraints outlined earlier [27, 28]. This innovative technology has transformed medical research, offering improved accuracy and efficiency in various applications related to personalized medicine [29].

Microfluidics has gained importance in cancer cell culture and drug discovery by enabling 3-D cell culture studies with both cancerous and non-cancerous cell lines (primary cells and stem cells) [30]. Microfluidic systems provide benefits like controlled blending, chemical gradients, reduced reagent use, constant perfusion, and precise control of pressure and shear stress on cells. [31]. Recent advancements in microfluidics have significantly contributed to the study of 3D spheroids by effectively addressing the limitations commonly associated with non-microfluidic techniques [32]. There are two types of microfluidic systems for 3D cell cultures: static microfluidic devices and continuous-flow microfluidic chips [33].

Polydimethylsiloxane (PDMS) has been increasingly explored for microfluidic devices and biological platforms for cell and tissue engineering applications. PDMS-based microfluidic devices display a precise micro-environment, minimal reagent requirements, and low cost of experiment [34]. PDMS is both optically transparent and gas-permeable, making it suitable for visualization [35]. The compact structure of the PDMS microfluidic device helps in designing microscale culture chambers [36]. However, the inherent hydrophobicity helps it to be an inert material in cell culture studies. For these reasons, PDMS became a popular polymer for applications ranging from MEMS to biomedical purposes [37].

Here, we have designed a high-throughput PDMS microwell device with hydrophobic and soft mechanical properties that can generate uniformly sized cancer spheroids. PDMS is non-toxic and biocompatible in many cellular studies. Due to its hydrophobic nature and contact angle greater than 100 degrees, PDMS can hold micro-volumes of liquids.

## Materials and methods

The PDMS (Sylgard^®^ 184, Dow Corning Korea, Seoul, Korea) was procured from Kevin Electrochem, Mumbai, India. AggreWell™ 400 and AggreWell™ 800 plates were purchased from STEMCELL Technologies, Vancouver, BC, Canada. Trichloro (1H,1H,2H,2H-perfluorooctyl) silane was purchased from SIGMA-ALDRICH, USA. Dulbecco’s Modified Eagle Medium High glucose (DMEM), Penicillin Streptomycin (Pen-strep), Paraformaldehyde (PFA), and Fetal Bovine Serum (FBS) were procured from Himedia, Mumbai, India. Phosphate buffer saline (pH 7), Collagen, was procured from Corning®, USA. Type 1 (Milli Q) water was used after two autoclaving cycles. All chemicals utilized were sterile and analytically pure and were used directly.

### Fabrication of molds and devices

### Fabrication of molds

The molds were created by replicating AggreWell™ plates with polydimethylsiloxane (PDMS). To prepare the mold, the AggreWell™ plate surfaces underwent silanization. The plates were positioned to avoid contact with the bottom, allowing the wells to remain exposed for effective silanization. Trichloro(1H,1H,2H,2H-perfluorooctyl) silane served as the silanizing agent. A few drops of this silane were placed on tissue paper and then put in a vacuum desiccator with the AggreWell™ plates overnight. A Sylgard 184 prepolymer was mixed with a curing agent in a 10:1 weight ratio, degassed, and poured over the silanized plates, where it cured for 2 hours at 70°C. After curing, the molds were removed from the AggreWell™ plates and utilized as master molds for microwell device fabrication.

### Fabrication of devices

The molds were silanized similarly to AggreWell™ plates. The pre-polymer resin mixture was degassed, then a thin layer was poured into a 150 × 25 mm polystyrene tissue culture dish and partially cured. Afterward, the silanized molds were placed inverted to ensure the correct embedding of microneedle structures. Following curing at 70°C for 2 hours in an oven, the solidified PDMS microwell devices were gently peeled from the molds.

*Figure 1* illustrates the detailed schematic of the step-by-step process for fabricating master molds and devices fabricated.

**Figure 1:**
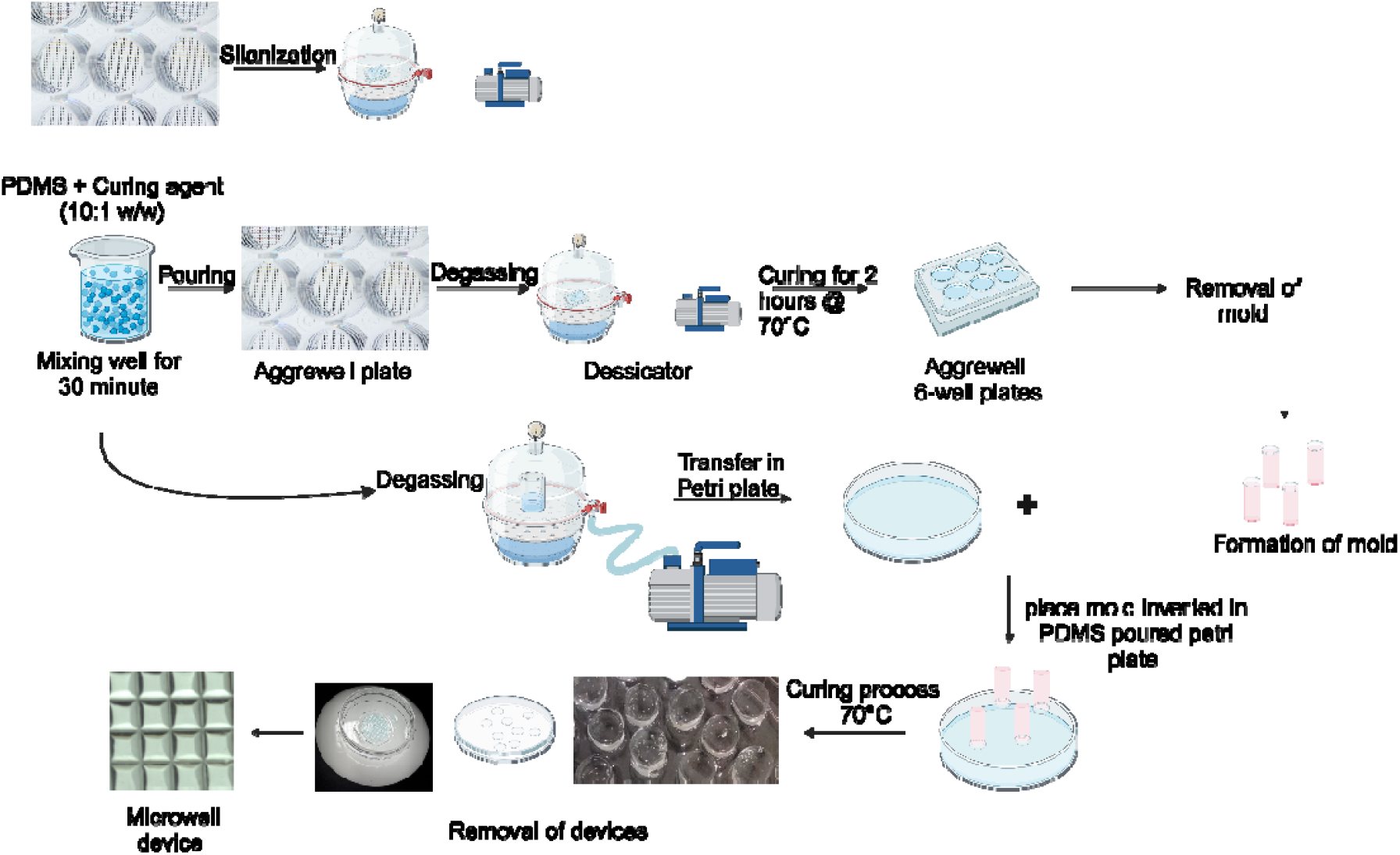
Schematic depicting the fabrication of molds and devices using the soft lithography technique

### Characterization of devices

#### Contact angle

Given that surface hydrophilicity and hydrophobicity were crucial factors in device fabrication, we measured the contact angle of the molds and devices using the Attension Theta Flex Optical Tensiometer. The analysis employed the sessile drop method, where a droplet of ultrapure Type 1 water was placed on the molds’ and devices’ surfaces. The contact angle was determined by fitting a tangent to the three-phase contact point at the intersection of the droplet surface and the PDMS surface. A digital video camera was used for visualization, and the contact angle was calculated with OneAttension software.

#### SEM imaging

Morphological characterization was conducted using Scanning Electron Microscopy (SEM) to analyze the formed microwells and generated spheroids. The PDMS devices were transversely sectioned to clarify the microwell cross-section and measure their dimensions. Before imaging, the spheroids were fixed in a 4% paraformaldehyde solution for 30 minutes, washed with PBS, and allowed to dry in a lyophilizer. The device samples and the spheroids were attached to SEM stubs with carbon tape and subsequently coated with platinum for 2 minutes to enhance conductivity and prevent sample charging. The prepared samples were then examined with SEM (Jeol-7600F).

#### Cell culture

To verify the fabricated device’s capability to generate spheroids, several cell lines were utilized. This study employed SH-SY5Y, HT-29, MCF-7, Caco-2, and MDA-MB-231 cell lines. SH-SY5Y is a human neuroblastoma cell line widely used in neuroscientific research. HT-29, derived from a White female patient, is a colorectal adenocarcinoma cell line commonly applied in cancer and toxicology studies. MCF-7 is a well-known human breast cancer cell line that has been extensively researched for over 45 years. MDA-MB-231, a triple-negative breast cancer (TNBC) cell line, is frequently used as a model for late-stage breast cancer.

All cell lines were cultured and maintained in Dulbecco’s Modified Eagle’s Medium (DMEM) with 10% fetal bovine serum and 1% antimycotic-antibiotic solution, incubated at 37°C in a humidified atmosphere with 5% COLJ.

#### Seeding and formation of spheroids

Cells were passaged when they reached about 80% confluency using 0.25% trypsin-EDTA. The device was positioned in a 60 mm Petri dish and pressed against the bottom for stability. Before seeding the cells, the device was rinsed with phosphate-buffered saline (PBS) to eliminate trapped air from the microwells. A pipette held at a 90° angle was used to seed the cells, ensuring the suspension was evenly distributed across the device. The seeded devices were incubated for 48-72 hours in a CO_2_ incubator, with media changes made regularly. During media exchange, the superficial medium was gently removed from the corner of the dish to avoid disturbing the cell aggregates, and fresh medium was introduced slowly from the same corner. To maintain humidity, PBS was added to the corners of the Petri dish, which was then covered with its lid and placed in the incubator. Figure 2 provides a stepwise illustration of cellular seeding and spheroid formation. After 48-72 hours, the formed spheroids were transferred into a collagen matrix, fixed, and stained with fluorescent dyes.

**Figure 2:**
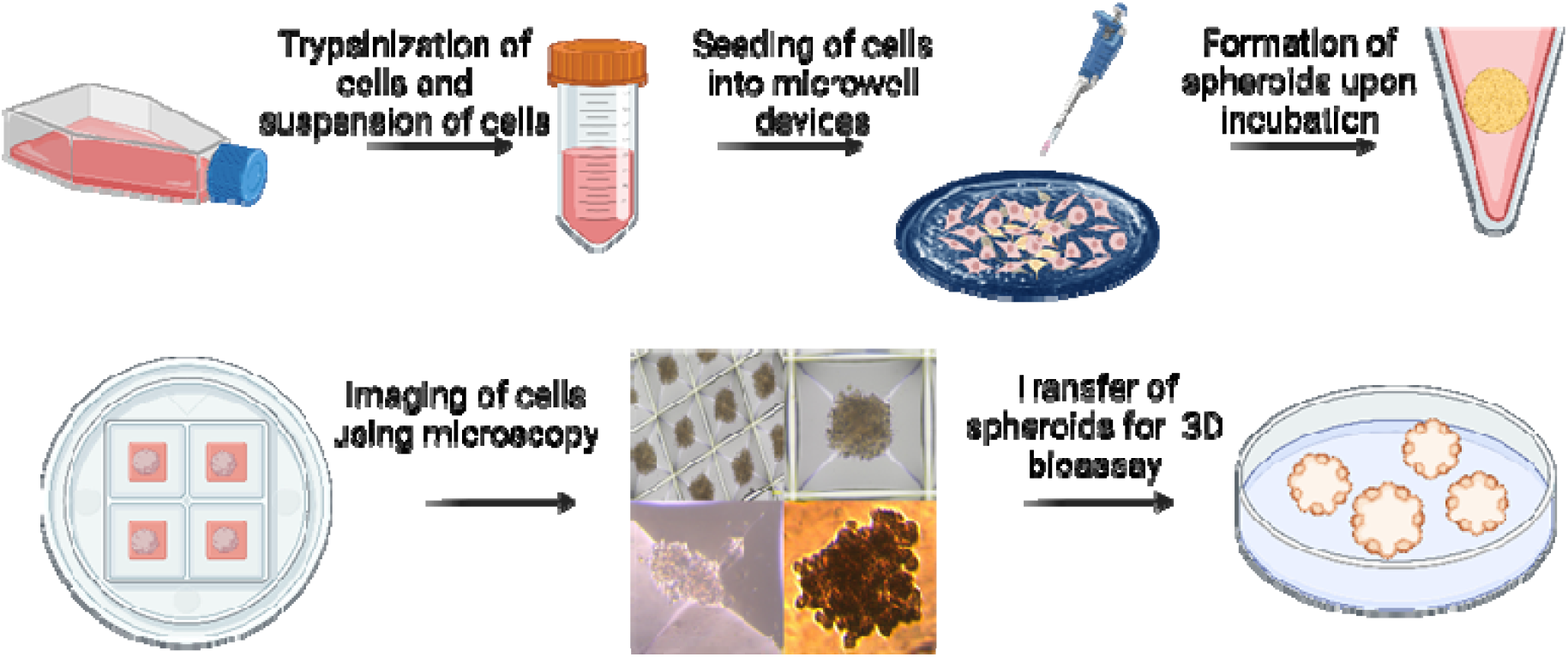
Schematic depicting the seeding of cells into microwell devices and the formation of spheroids.

For nuclear and viability staining, the fixed spheroids were incubated with Hoechst 33342, a fluorescent DNA-binding dye, at a final concentration of 1 μg/mL for 20 min at room temperature. Propidium iodide (PI) stock solution (1 mg/mL) was diluted at a 1:3000 ratio in 1× phosphate-buffered saline (PBS) and added to the spheroids to allow selective staining of permeabilized dead cells. After staining, the spheroids were washed three times with 1× PBS, and coverslips were mounted onto glass slides for imaging.

#### Image processing and Statistical analysis

Fluorescence images of stained spheroids were acquired using a confocal microscope with appropriate excitation and emission settings. Image processing and quantitative analysis were performed using ImageJ (version [insert version number]), where spheroid size, circularity, and fluorescence intensity were analysed. Image pre-processing steps included background subtraction, contrast enhancement, and threshold adjustment to optimize visualization and quantification.

Data plotting and statistical analyses were conducted using GraphPad Prism (version [insert version number]). Statistical significance was determined using appropriate tests, such as Student’s t-test or one-way ANOVA, based on the experimental design. All data were expressed as mean ± standard deviation (SD) unless stated otherwise. A p-value < 0.05 was considered statistically significant.

## Results and Discussion

### Molds & Devices

PDMS molds were fabricated using standard soft lithography techniques. For mold fabrication, AggreWell™ 24-well plates with microwell sizes of 400 μm and were used as templates to replicate the topographical features of the PDMS microwell devices. Silanization of the AggreWell™ plates was performed to ensure easy peeling of molds from the well plates. Successful silanization was confirmed by the visible transition of the AggreWell™ plates from a transparent to a hazy appearance.

Molds with a 13 mm diameter, containing microneedles 400 μm in height, were fabricated and subsequently used for the production of PDMS microwell devices. These molds served as negative replicas of the AggreWell™ plates. Both the AggreWell™ plates and the fabricated molds were silanized before device generation. The silanization process effectively reduced surface adhesion between the mold surface and the device fabricated, enabling easy peeling of PDMS devices from the mold surfaces. The resulting PDMS microwell devices were flexible, mechanically robust, and exhibited uniformly distributed wells that remained structurally intact upon demolding. (Figure 3).

**Figure 3:**
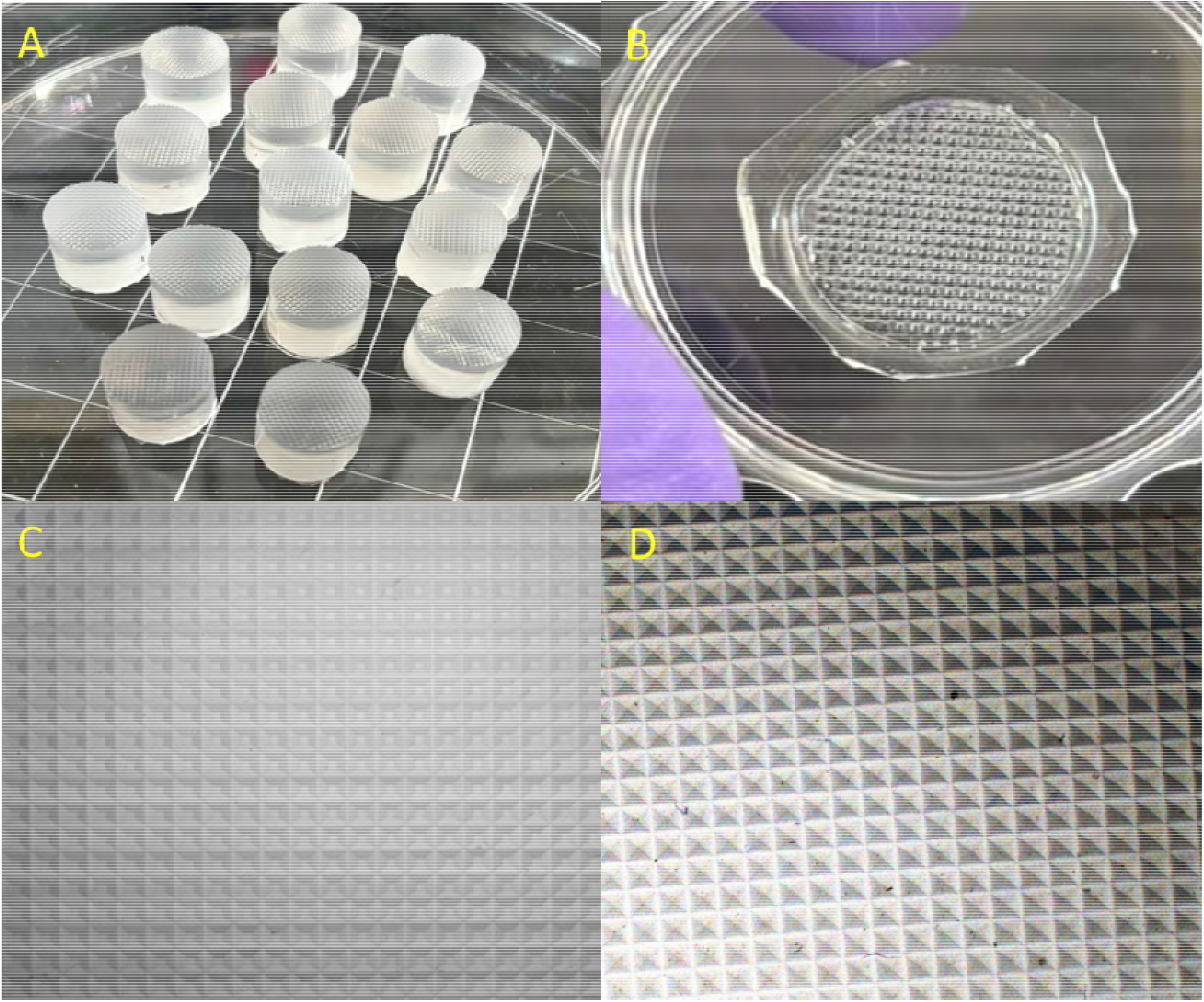
Images of fabricated PDMS molds and devices using soft lithography method. A) PDMS molds, B) PDMS devices C) PDMS devices imaged using microscope d) PDMS molds imaged using microscope.

Conversely, unsilanized molds led to the creation of defective devices with uneven wells, tearing, and compromised structural integrity, resulting in peeling from the mold (Figure 4). The PDMS device bonded tightly to the unsilanized mold, making detachment difficult. Figure 4 showcases an example of a device fused to the mold, emphasizing the importance of silanization for producing high-quality PDMS microwells.

**Figure 4:**
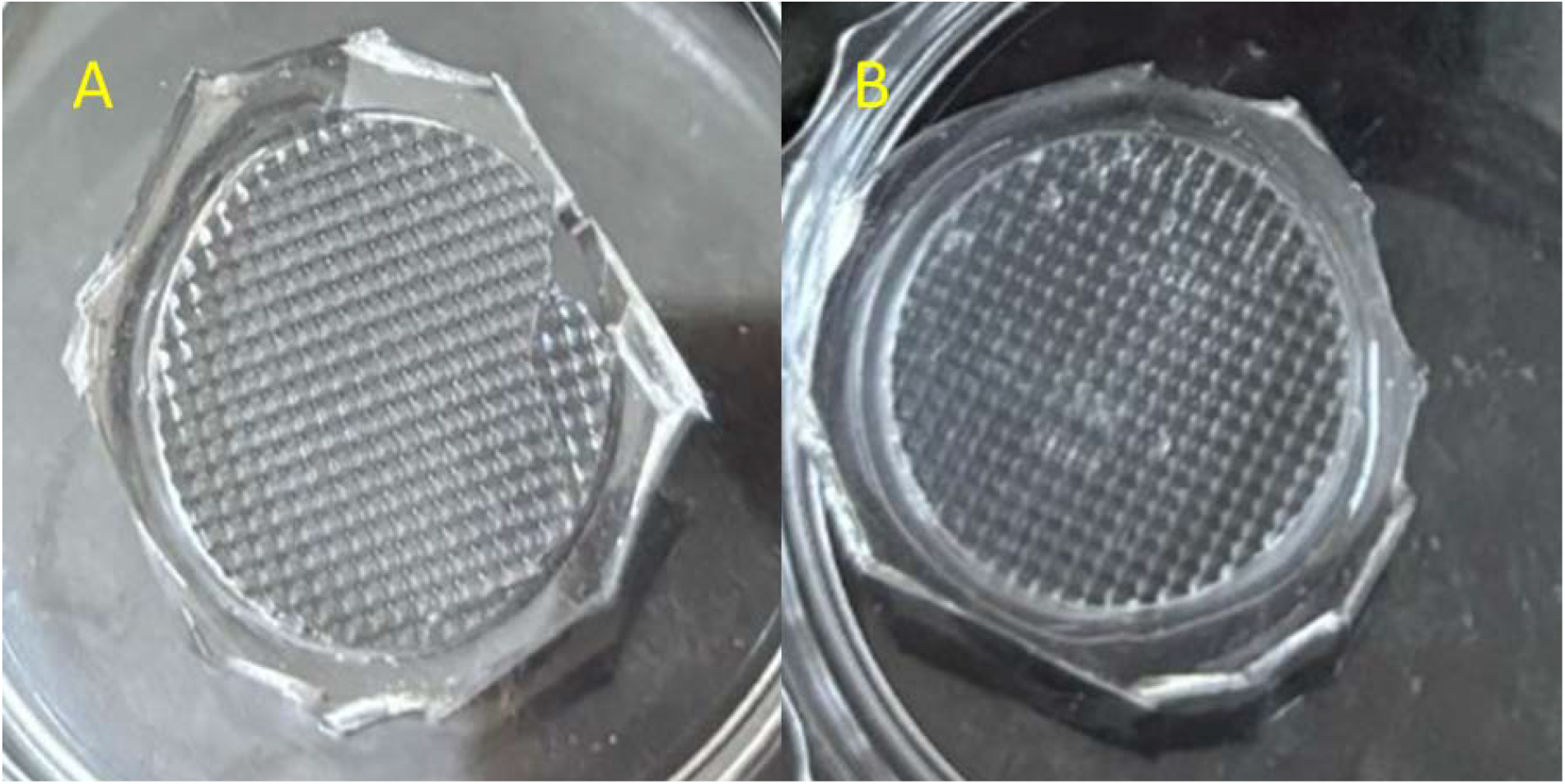
Damage caused to the device during peeling from unsilanized molds. A&B) Image of device torn and structure damage occurred during peeling

**Figure 5:**
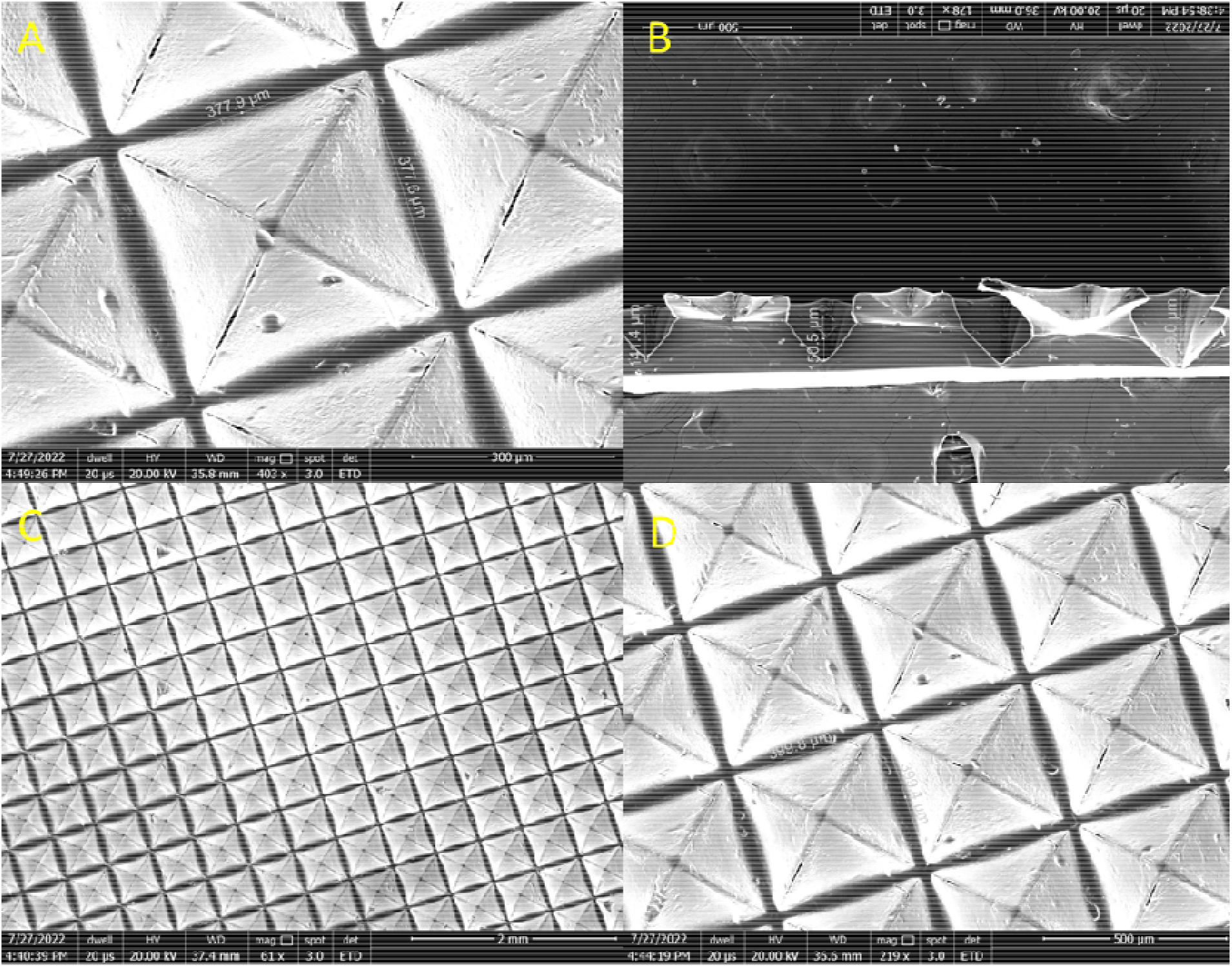
Scanning Electron Microscope imaging of PDMS microwell devices. Vertical view and cross-section of devices

### Characterization of microwell devices

The engineered PDMS microwell devices possess optical transparency and the capacity to hold liquid volumes up to 1 ml. These devices demonstrate flexibility and chemical inertness, rendering them appropriate for a variety of cell culture applications. Given their compact design, featuring a diameter of 13 mm, the microwell devices can be positioned within any standard culture dish or well plate possessing a diameter greater than 14 mm, thereby ensuring compatibility with multiple experimental setups.

### Morphology of the devices

Scanning electron microscopy (SEM) was employed to analyse the morphological characteristics of the fabricated PDMS microwell devices. The wells exhibited an inverted hollow prism geometry with dimensions of approximately 400 × 400 μm. To determine the depth of the microwells, SEM imaging was performed on cross-sectional samples of the device. The depth of the microwells was measured to be 146 ± 6 μm, as depicted in the SEM images. The SEM images of the devices further revealed that the microwell walls were smooth and flat, devoid of any surface texture. This structural characteristic is critical for spheroid formation, as the absence of surface roughness prevents cellular adhesion to the device, thereby encouraging cell aggregation within the wells. Consequently, the cells formed spheroids by adhering to one another rather than attaching to the microwell surface.

### Contact angle of microwell molds and devices

One of the primary challenges in fabricating microfluidic devices is the ability to modulate surface properties, such as hydrophilicity, to suit cellular applications. To assess these properties, the contact angles of PDMS molds before and after silanization and microwell devices were measured. (Table 1) presents the sessile drop method measurements, detailing the contact angles observed under different conditions. Prior to silanization, the contact angle of PDMS molds was measured to be 130 ± 2° (Figure 6A), indicating inherent hydrophobicity. Following silanization, the contact angle increased to 150 ± 2° (Figure 6B), confirming the successful surface modification. This 20° increase can be attributed to the deposition of the silane layer, which further enhanced surface hydrophobicity. Similarly, the contact angle of the fabricated PDMS microwell devices was measured to be 130 ± 2°, demonstrating that the devices retained their hydrophobic nature post-fabrication. The intrinsic hydrophobicity of the PDMS microwell devices played a crucial role in spheroid formation. The hydrophobic surface repelled culture media droplets, preventing cell adhesion to the device surface. As a result, the suspended cells within the microwells aggregated, facilitating the formation of uniform spheroids. Figure 6 C& D illustrates the contact angle of 113 ± 2° for both indicating the hydrophobicity of devices which is the reason behind non-adhering of cells leading to formation of spheroids.

**Table 1:** Variation in the contact angle of different surfaces before and after surface modification and contact angle measurement of microwell devices (800 μm) & (400 μm)

|  |  |  |
| --- | --- | --- |
| 1 | Master mold | $130 \pm 2^\circ$ |
| 2 | Master mold after salinization | $150 \pm 2^\circ$ |
| 3 | Microwell Device (800 $\mu\text{m}$) | $113 \pm 2^\circ$ |
| 4 | Microwell Device (400 $\mu\text{m}$) | $113 \pm 2^\circ$ |

**Figure 6:**
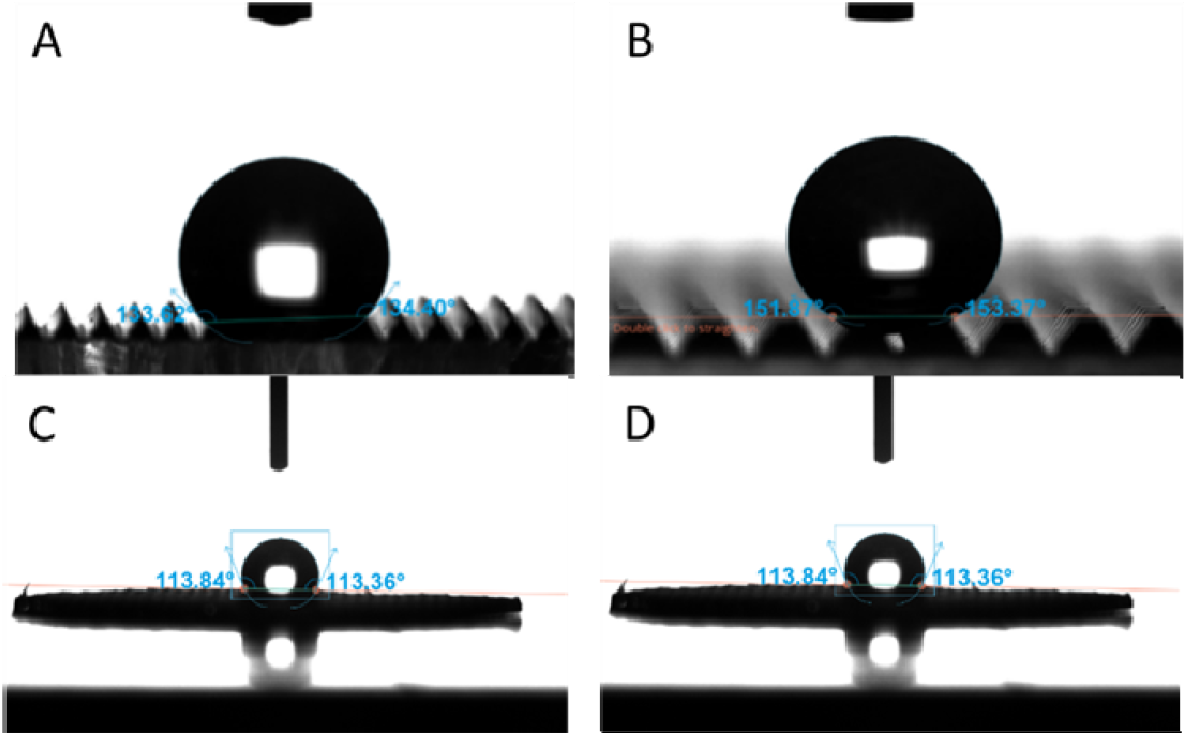
Contact angle measurement of PDMS molds to show that the silanization resulted inthe conversion of PDMS molds surface from hydrophobic to superhydrophobic

**Figure 7:**
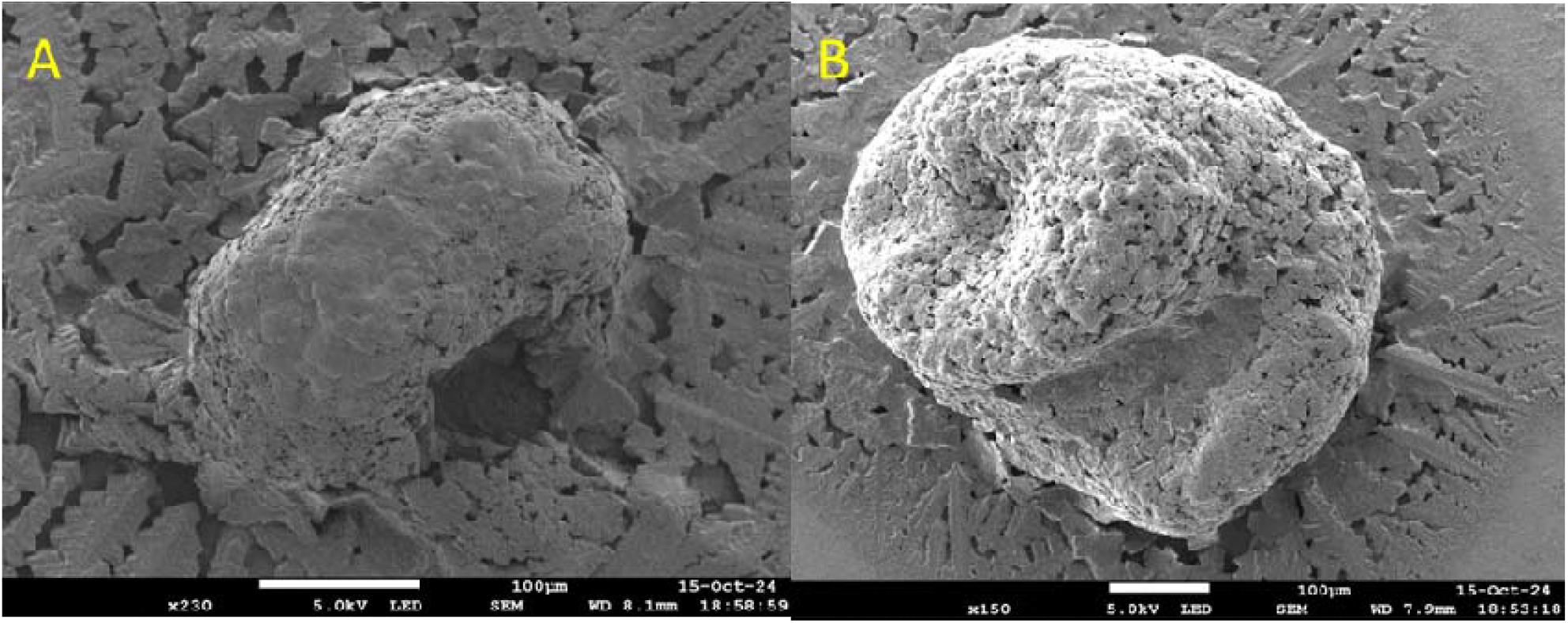
SEM image of A) Caco2 spheroid formed using 400 μm device, B) HT-29 spheroid formed using 800 μm device

### Formation of spheroids in the microwell device

Before seeding the cells, the PDMS microwell devices were inverted and placed in a beaker with autoclaved Milli-Q water, then sonicated at room temperature to remove trapped air bubbles from the microwells. Subsequently, the devices were dried and sterilized through autoclaving, preparing them for cell culture experiments. Each PDMS microwell device featured 1,200 microwells per well, designed for 400 μm-sized microwells. A fixed cell counts of 100 cells per microwell (0.12 million cells per mL in 400 μm devices) and 3000 cells per microwell (0.9 million cells per mL in 800 μm devices) was maintained across all cell lines in the fabricated device. Aggregation of cells was visible within 24 hours post-seeding, with the development of distinct three-dimensional (3D) spheroid structures occurring between 48 and 72 hours. The duration for spheroid formation varied among the different cell lines. The resulting spheroids were well-formed, tightly packed, and remained intact during media exchange, indicative of robust cell-cell interactions. Microscopy images verified that the cells did not adhere to the microwell surfaces but rather aggregated together to create spheroids.

The spheroid sizes ranged from approximately 200 μm to 700 μm, depending on the cell line and type of microwell device used (400 μm or 800 μm). Notably, spheroids formed by the MDA-MB-231 triple-negative breast cancer (TNBC) cell line exhibited a more defined spherical morphology, consistent with its aggressive and metastatic nature. A single 400 μm microwell device facilitated the formation of approximately up to 1,200 homogeneous spheroids in a single experiment.

The spheroids were freely suspended in the culture medium within the microwells, which facilitated easy harvesting. By gently dispensing fresh medium, the spheroids were dislodged from the wells, allowing for their collection through aspiration using a serological pipette. The harvested spheroids were then placed onto coverslips that had been pre-coated with a collagen matrix and incubated at 37°C in a 5% CO□ incubator for one hour. After the transfer, some spheroids underwent fluorescence staining for imaging purposes. Figure 8 illustrates representative confocal images of spheroids from different cell lines. The images obtained were quantified for their diameter and circularity, with the results presented in Table 2. The findings indicate that all selected cell lines produced compact, spherical, and intact spheroids. Of the five cell lines, the MDA-MB-231 line produced perfectly spherical spheroids with a circularity of 0.9, while Caco2 formed spheroids that were not entirely intact. The spheroids formed within a timeframe of 48-72 hours; the longest time, 72 hours, was observed for the KB, MDA-MB-239, and HT-29 cell lines, whereas MCF-7 and Caco2 achieved spheroid formation in the shortest interval of 48 hours. This variation may be attributed to the rapid formation of gap junctions between the cells. Table 2 summarizes the area, circularity, and formation time of the spheroids.

**Table 2:** Variations in area of spheroid and circularity for different spheroids are detailed along with the time taken for formation

| Cell line | Area of spheroid<br>(Sq. Micron) | Circularity of<br>spheroid | Time taken for<br>formation (Hrs) |
| --- | --- | --- | --- |
| **KB 31** | 101927.7 (± 1061.4) | 0.858 (± 0.02) | 72 h |
| **MCF 7** | 317940 (± 3156.4) | 0.782 (± 0.05) | 48 h |
| **MDA MB 231** | 98843 (± 2158.9) | 0.925667 (± 0.004) | 72 h |
| **Caco<sub>2</sub>** | 197846.7 (± 3046.7) | 0.749667 (± 0.03) | 48 h |
| **HT** | 258063.7 (± 2809.4) | 0.886667 (± 0.02) | 72 h |

**Figure 8:**
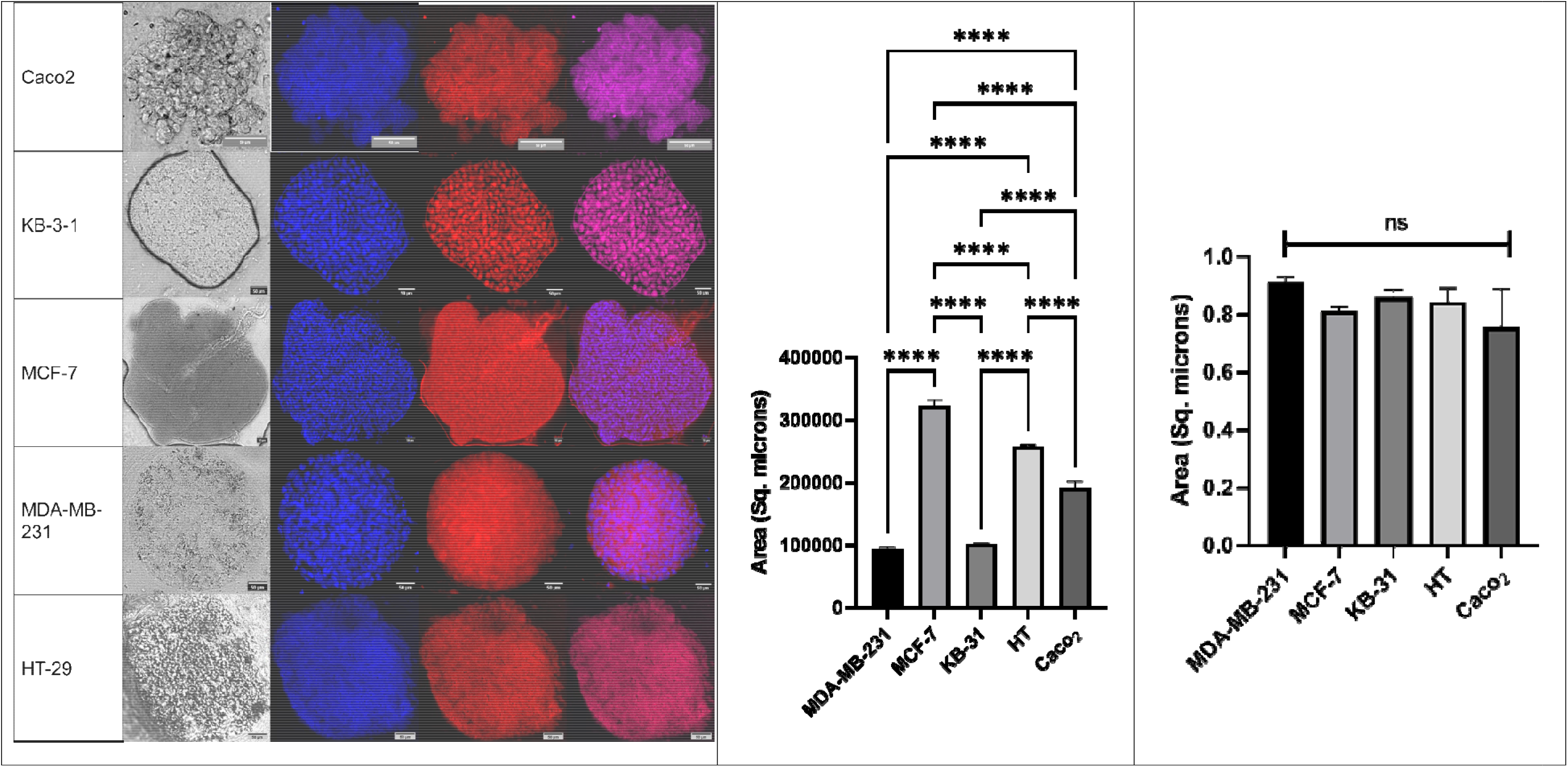
(A) confocal images of the spheroids formed and stained with DAPI (Blue), PI (Red), Bright field (Grey) and Merged channels. (B) Quantification data of Area of spheroids generated. (C) circularity data of spheroids generated.

### Cellular compatibility of the device

The media extracts imbibed with the fabricated devices were subjected to the cells seeded in 96 well plates and the viability of the cells was estimated (Figure 9) to be 98.7 % (± 1.0%), 101.6% (± 2.78%) and 99.6% (± 1.23%) viability of cells were calculated for the cells subjected to media incubated with devices for 24hrs, 48hrs and 72 hrs respectively. The viability percentages obtained from the cells treated with device-imbibed media resulted in no significant difference when compared to the viability of the control. It can be interpreted that the devices are completely bio-compatible and the device materials or the devices do not alter any cellular viability and cellular properties. The devices only facilitated the aggregation of cells onto each other, resulting in the formation of spheroids.

**Figure 9:**
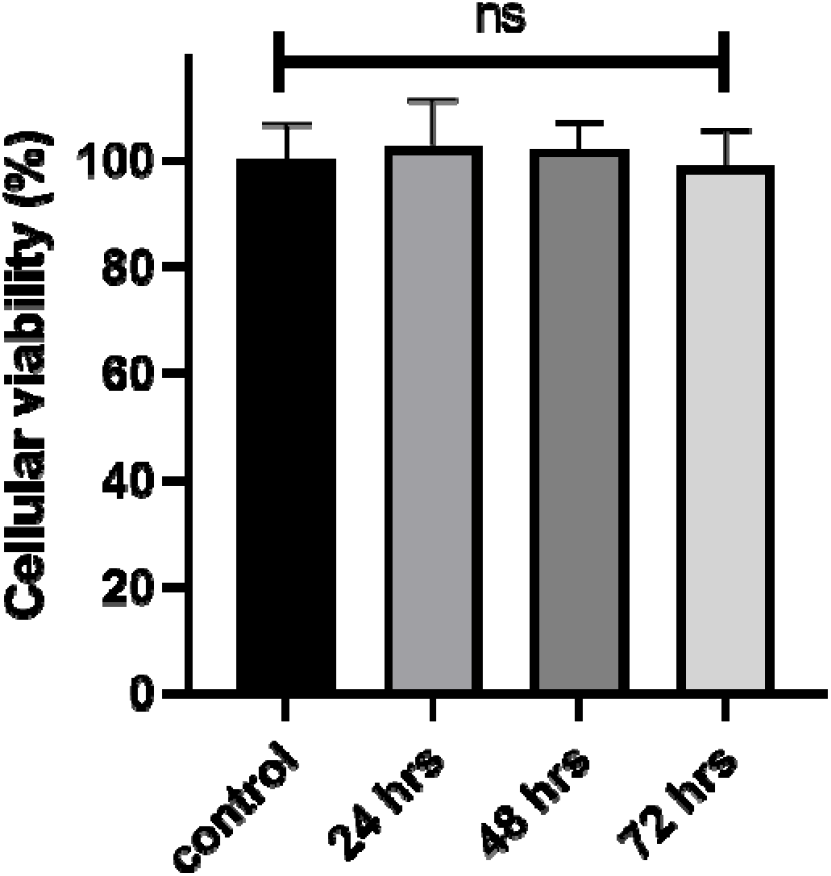
Graphical representation of cell viability data in the presence of PDMS microwell devices for 24, 48, 72 hours, respectively

## Conclusion

We introduce a microfluidic device engineered for the large-scale generation and in situ analysis of homogeneous cell spheroids. The 3D cell aggregates created by this device provide a more “tissue-mimetic” model when compared to conventional 2D monolayers. This paper underscores the utility of microfluidic technology in spheroid formation and culture, elaborating on its potential benefits. We expect this system to enhance high-throughput drug testing on both cell line spheroids and patient-derived samples. Moving forward, the development of tissue models that can offer more accurate biomimetic in-vivo representations is essential, as this will result in extensive screening data. We have engineered a polydimethylsiloxane (PDMS)-based microwell device to generate uniformly sized spheroids from cells suspended in media. Our design is compatible with standard Petri dishes and well plates, ensuring straightforward integration into current cell culture practices. The PDMS microwell device includes microwells with a 400-micron diameter, and we believe our culture system presents significant potential for various applications in fields such as tissue engineering and drug screening.

